# InfluProto: Predicting Cross-Species Transmission of Influenza A Virus via Prototype Learning

**DOI:** 10.64898/2026.10.07.757480

**Authors:** Zongyi Geng, Lun Li, Rongye Ye, Yi Wang, Shuhui Song

## Abstract

Zoonotic influenza A viruses constitute a persistent threat to global public health, yet quantitative approaches for assessing their cross-species transmission potential remain limited. Here, we present InfluProto, a deep learning model that introduces prototype learning into host prediction for influenza A virus, conceptualizing host adaptation as a continuous process rather than a discrete outcome. InfluProto embeds viral sequences into a continuous representation space anchored by human, swine, and avian prototypes, using cosine distances as a quantitative measure of host adaptation. InfluProto accurately discriminated among the three host classes, achieving recall and precision exceeding 0.97, and substantially outperformed conventional classification models*—*particularly on challenging zoonotic strains, including the 2009 pandemic H1N1 (pdm09) lineage. Beyond host classification, the learned representations captured biologically meaningful host-adaptive signatures; notably, pdm09 viruses occupied an intermediate position between human and swine clusters, consistent with their evolutionary origin. Analysis of simulated reassortment further revealed that the introduction of human-origin segments into swine influenza viruses reduced their distance to the human prototype, indicating that InfluProto can detect reassortment-associated shifts in host adaptation. To facilitate broad public access, we have developed an interactive web service for InfluProto. Collectively, our work establishes a continuous framework for characterizing host adaptation and evaluating zoonotic risk in influenza A viruses, with potential implications for pandemic surveillance and preparedness.

## Introduction

Zoonotic diseases account for an estimated 70–80% of emerging infectious diseases and pose a persistent threat to global public health[1, 2]. Among zoonotic pathogens, influenza A virus (IAV) is of particular concern because of its broad host range[3] and remarkably rapid evolution. IAV evolves rapidly through two principal mechanisms: antigenic drift and antigenic shift. Antigenic drift involves the gradual accumulation of mutations, particularly in genes encoding surface glycoproteins. Antigenic shift, by contrast, arise from the reassortment of viral gene segments and can lead to substantial changes in viral phenotypes, including host tropism[4]. Both evolutionary processes have repeatedly contributed to the emergence of pandemic strains, such as those causing the 1918 H1N1 pandemic and the 2009 H1N1 (pdm09) pandemic, as well as the emergence of H7N9 avian influenza in humans[2]. Such events emphasize the importance of host prediction and risk assessment for zoonotic strains in order to mitigate the severity of future outbreaks[2].

Traditional wet-laboratory methods remain the gold standard for assessing viral host tropism. However, they are time-consuming, costly, and generally low-throughput, which limits their application to large-scale screening of the vast number of viral strains generated through global influenza surveillance. In addition, experimental assessment of host tropism often relies on animal- or cell-based systems, which may be confounded by experimental and environmental factors[5, 6].

Beyond experimental assessment, viral host tropism can be inferred from sequence information using two main approaches. One approach infers viral host tropism based on prior biological knowledge, particularly known host-adaptive mutations[7]. However, this approach is inherently constrained by the breadth and accuracy of prior knowledge. It is largely limited to previously characterized sites and may therefore overlook novel or previously undescribed determinants of host specificity. Another approach infers viral host tropism directly from sequence information using deep learning, which has been increasingly applied to influenza virus research[8–10]. Early models were often developed using data from multiple virus families, which limited their ability to capture influenza-specific evolutionary patterns[11, 12]. Subsequent influenza-specific models were typically developed using individual gene segments. However, these models often perform poorly on zoonotic strains and may generate conflicting predictions across different segments[13, 14]. More recent models leverage full-genome inputs, yet its performance on zoonotic viruses remains limited, and the risk indicators depend heavily on the composition of reference datasets[15]. Underlying these limitations is a deeper conceptual issue that host adaptation in viruses is inherently a continuous evolutionary process, not a discrete classification event. During intermediate evolutionary states, a virus may retain the ability to infect multiple host species simultaneously, which makes it difficult to classify. As a result, standard classification-based models are fundamentally misaligned with this biological reality.

To address this gap, we developed InfluProto, a prototype-learning framework that represents influenza A virus host adaptation in a continuous embedding space[16, 17]. The model learns representative prototypes for human-, swine-, and avian-hosted viruses, and characterizes viral sequences according to their relative distances to these prototypes, rather than assigning them to a single host category. This representation provides a continuous profile of host adaptation, allowing intermediate states to be characterized without forcing viruses into discrete host categories.

InfluProto is trained on whole genome sequences of host-restricted influenza A viruses from GISAID[18] and evaluated on three independent test datasets. Notably, the model outperformed conventional classification models, particularly on a zoonotic dataset of the pdm09 lineage, a well-known zoonotic lineage[19]. Beyond predictive performance, the learned representation also shows biologically meaningful patterns. Specifically, attention is concentrated on sites associated with host adaptation. Meanwhile, distances to the three prototypes quantitatively reflect variation in host adaptation across hosts and clades. An interactive web server was developed to facilitate sequence-based assessment and exploration of prototype-space relationships (https://ngdc.cncb.ac.cn/bit/influProto). Collectively, this work establishes a continuous and interpretable framework for zoonotic risk evaluation in influenza viruses and provides algorithmic infrastructure ready for prospective surveillance of future cross-species transmission events.

## Results

### Architecture of InfluProto: Prototype Learning for Continuous Host Adaptation

To capture the continuous nature of host adaptation, InfluProto employs prototype learning to map viral sequences into a continuous representation space defined by host-specific prototypes. Within this framework, host adaptation is characterized by the relative proximity of each viral sequence to the human, swine, and avian prototypes.

Specifically, InfluProto takes whole-genome sequences of influenza A viruses as input and processes them through a three-stage architecture (Figure 1). First, a MEGA encoder[20] extracts per-position features from the input sequence. This encoder was selected for its demonstrated efficacy in modeling long genomic sequences, making it particularly suitable for whole-genome analysis of influenza A viruses[15]. Second, an attention pooling layer assigns a learnable weight to each nucleotide position and aggregates position-specific features into a single 128-dimensional embedding vector. Third, instead of employing a conventional classification layer, the model computes the cosine distance between the embedding vector and three learnable prototype vectors, each representing viruses of human, swine, or avian origin. A smaller distance to a given prototype indicates closer alignment with that host and, consequently, a higher predicted likelihood of infecting that host. The model was optimized using a cross-entropy loss function applied over the negative distances (detailed in Methods).

**Figure 1.**
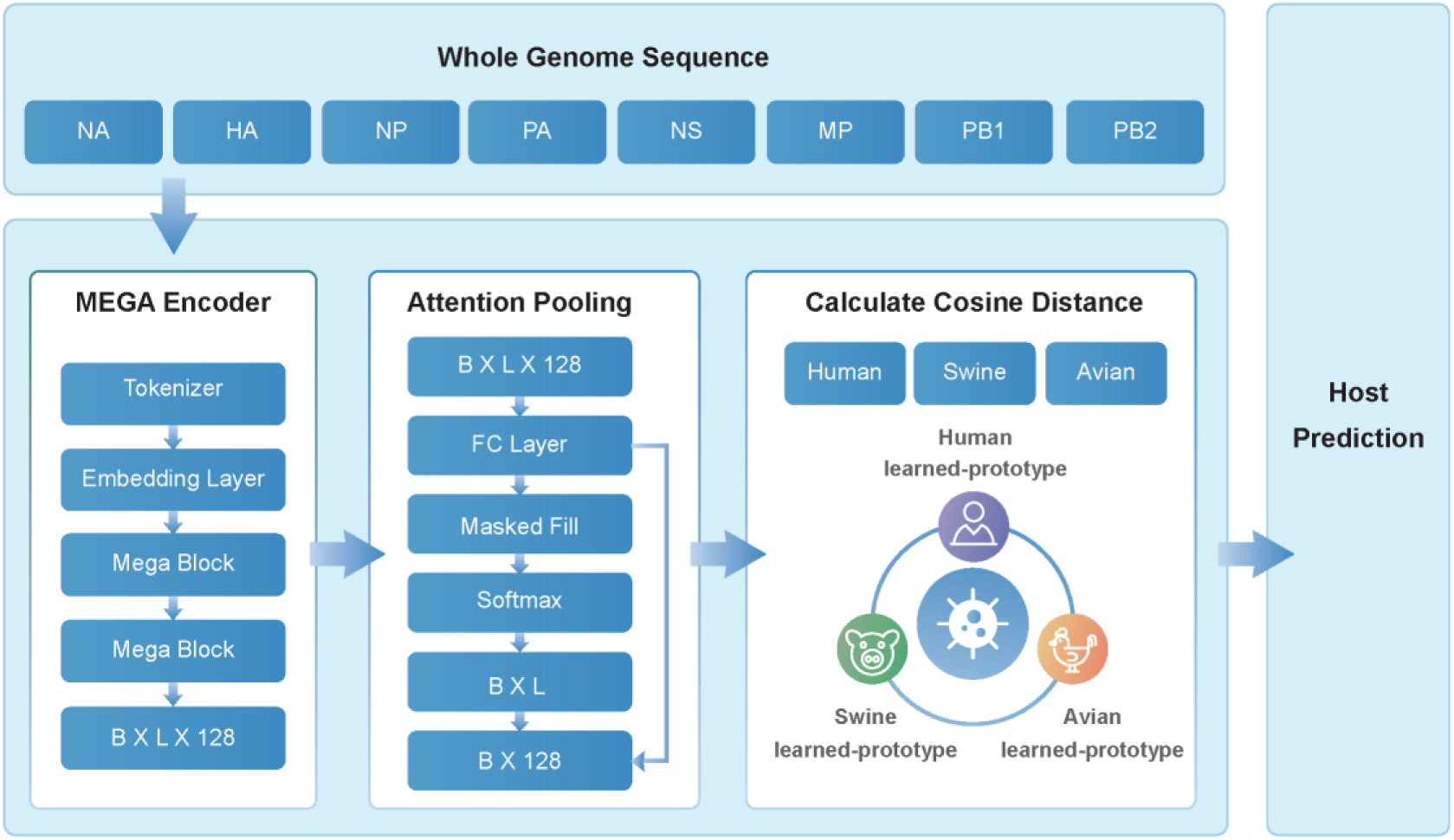
Architecture of InfluProto. The model takes the eight concatenated genome segments of an influenza A virus as input. A MEGA encoder first extracts per-position features, which are then aggregateed by an attention pooling layer into a single 128-dimensional embedding vector *E*(*v*), where *B* denotes the batch size, and *L* denotes sequence length. Three learnable prototype vectors represent the prototypical feature positions of viruses associated with human, swine, and avian hosts, respectively. Host prediction is achieved by computing the cosine distance between *E*(*v*) and each prototype; the prototype with the smallest distance determines the predicted host.

The model was trained on a *host-restricted dataset* comprising approximately 30,000 influenza A virus sequences, partitioned into training, validation, and test sets at an 8:1:1 ratio. To avoid ambiguous supervisory signals, sequences with documented evidence of cross-species transmission were excluded from model training. To further evaluate the model’s generalizability to viruses with zoonotic potential, we constructed three independent evaluation datasets: the *2009 pandemic H1N1 (pdm09) lineage*, a *zoonotic avian dataset*, and a *zoonotic swine dataset* (detailed in Methods and Supplementary Figure 1).

To evaluate the contributions of prototype learning and attention-based pooling, we further constructed two baseline classification models using the same MEGA encoder backbone. Baseline 1 replaces the prototype layer with a conventional linear classifier while retaining attention pooling. Building on this, baseline 2 further replaces attention pooling with mean pooling. Together, these baselines represent the standard discrete classification paradigm, providing a direct comparison to the continuous framework proposed in InfluProto.

### InfluProto Accurately Discriminates among Human, Swine, and Avian Hosts

We first evaluated the host classification performance of InfluProto on the held-out test dataset, comprising 10% of the approximately 30,000 sequences in the *host-restricted dataset*. InfluProto achieved recall and precision exceeding 0.97 for all three host classes — human, swine, and avian (Figure 2a, 2b), with PR-AUC values exceeding 0.99 across all three classes (Figure 2c). Stratified analysis by viral subtype further confirmed the model’s robustness: prediction accuracies were consistently above 0.95 for the five most prevalent subtypes in the test dataset (Supplementary Figure 2a).

**Figure 2.**
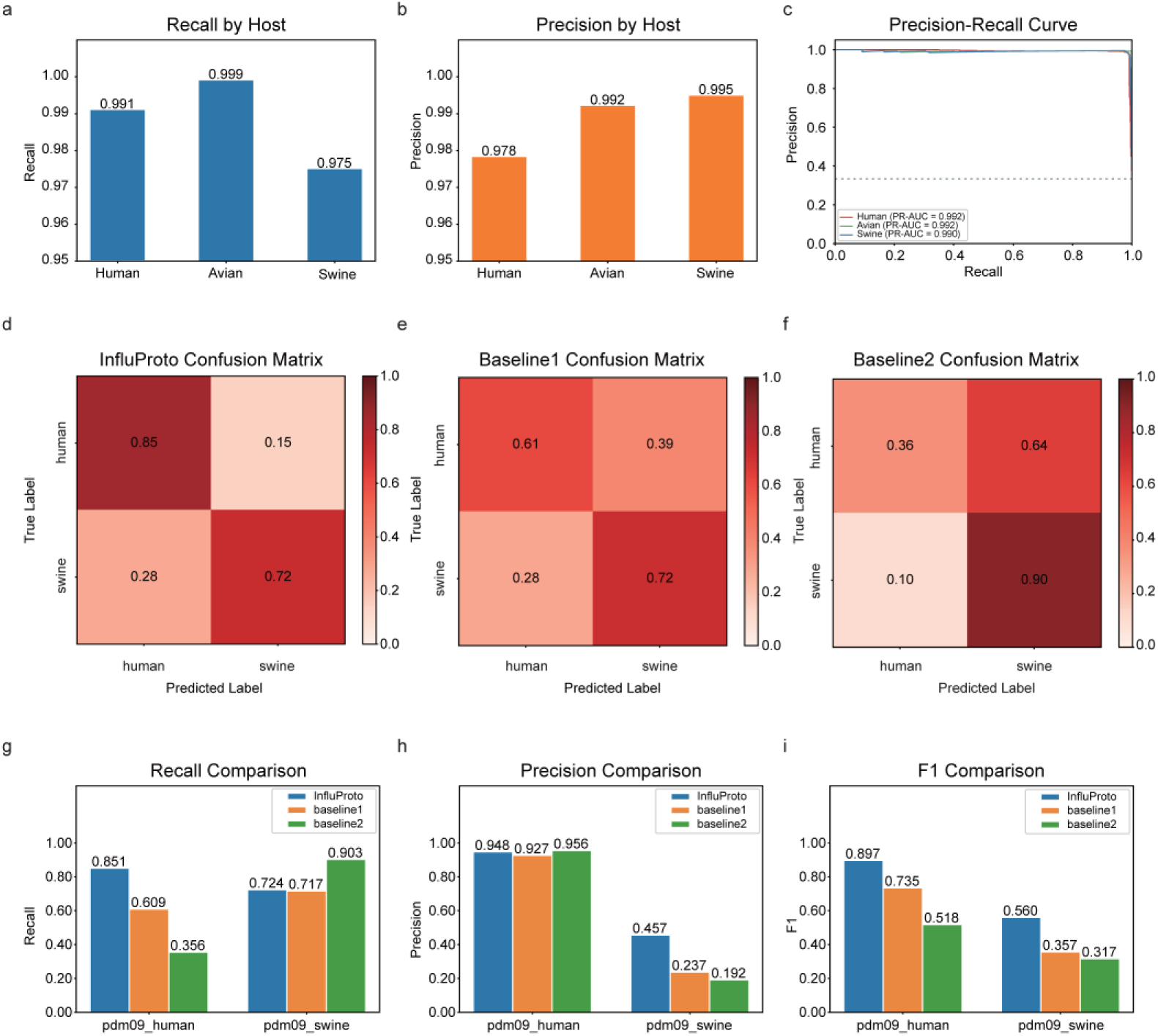
Prediction performance of InfluProto. **a–c**, Host prediction on the host-restricted test dataset. **a**, Recall across human, avian and swine hosts. **b**, Precision across human, avian, and swine hosts. **c**, PR-AUC curves for human avian and swine hosts. **d–f**, Comparison of InfluProto and the classification baseline models on the *pdm09 lineage dataset*. **d**, Recall for human-hosted and swine-hosted pdm09 sequences under each model. **e**, Precision for human-hosted and swine-hosted pdm09 sequences under each model. **f**, F1 score for human-hosted and swine-hosted pdm09 sequences under each model. **g–i,** Confusion matrices on the *pdm09 lineage dataset* for InfluProto (**g**), baseline 1 (**h**), and baseline 2 (**i**). Rows represent the true sampling host; columns represent the predicted host.

Having established strong performance on host-restricted sequences, we next assessed InfluProto’s ability to characterize host adaptation in zoonotic viruses by evaluating it on the *pdm09 lineage dataset*, which was entirely excluded from model training. Because pdm09 viruses infect both humans and swine, we computed recall and precision separately for each host and benchmarked InfluProto against the two baseline classification models. In this challenging zoonotic setting, InfluProto substantially outperformed both baselines. The baseline models tended to classify the majority of pdm09 sequences as swine regardless of their actual sampling host (Figure 2d, 2e, 2f), reflecting the known swine origin of the pdm09 lineage. Although this bias conferred higher recall on swine-hosted sequences, it resulted in markedly lower precision for that class, indicating a high false-positive rate (Figure 2g, 2h). InfluProto, in contrast, was more sensitive to the host-specific signals retained in human-isolated pdm09 sequences, enabling it to discriminate between human- and swine-origin isolates within this dual-host lineage. Specifically, InfluProto achieved roughly 25 and 50 percentage points higher recall than the two baseline models, respectively, while maintaining comparable precision (Figure 2g, 2h). The F1 score and PR-AUC (Figure 2i, Supplementary Figure 2b), which jointly balance recall and precision, consistently confirmed InfluProto’s overall superiority on the *pdm09 lineage dataset*. This advantage is particularly meaningful, as accurately identifying the human-infecting potential of zoonotic strains is precisely the capability of practical interest.

We further extended this evaluation to the *zoonotic swine dataset* and the *zoonotic avian dataset*, and the results were fully consistent with those observed for the *pdm09 lineage* (Supplementary Figure 2c, 2d), reinforcing InfluProto’s generalizable advantage in detecting zoonotic potential across diverse viral backgrounds.

### InfluProto Captures Biologically Meaningful Host Adaptation Signatures

To investigate whether the representations learned by InfluProto capture biologically meaningful patterns of host adaptation, we analyzed both its embedding space and attention patterns. The 128-dimensional viral embeddings generated by the attention pooling layer were projected into a two-dimensional space using t-SNE for visualization. In the embedding space, the host-restricted test dataset formed distinct clusters for human, swine, and avian viruses, with partial overlap between the human and swine clusters, reflecting the recognized role of swine as a mixing vessel for influenza viruses (Figure 3a).

**Figure 3.**
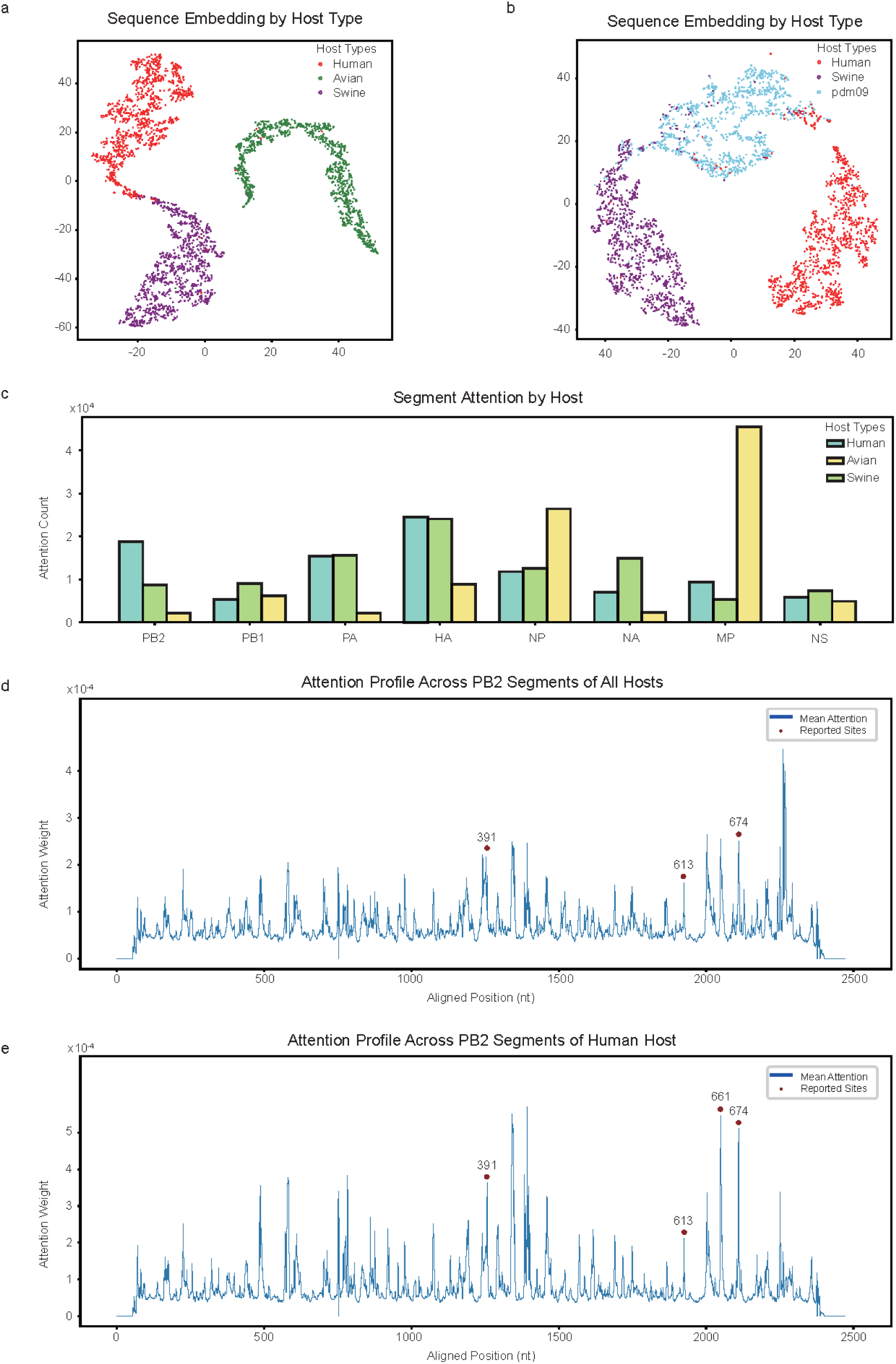
Interpretability analysis of InfluProto. **a**, t-SNE projection of the 128-dimensional embeddings of 3,000 host-restricted test dataset sequences, colored by host. **b**, t-SNE projection of 1,000 pdm09 sequences overlaid with human and swine host-restricted sequences. pdm09 embeddings fall between the human and swine clusters. **c**, Distribution of the top 100 most-attended positions across the eight genome segments, stratified by host. **d**, Per-position mean attention weights on the PB2 segment in all hosts, mapped to a reference sequence. **e**, Per-position mean attention weights on the PB2 segment in human derived sequences. Position 661, a known host-range determinant, receives substantially higher attention in human-hosted PB2 sequences than the global average.

We next projected embeddings of 1,000 randomly sampled pdm09 sequences, together with those of the human- and swine-hosted sequences from the host-restricted test dataset, into the same two-dimensional t-SNE space. The pdm09 embeddings occupied an intermediate position between the human and swine clusters (Figure 3b), reflecting the established dual-host nature of this zoonotic lineage. Similarly, the sequences from the *zoonotic avian dataset* fell between host-restricted human and avian clusters (Supplementary Figure 3a). When all of the above datasets were projected together into the same t-SNE space, the resulting global landscape recapitulated the patterns observed in the individual projections: host-restricted viruses formed distinct clusters, while zoonotic viruses occupied intermediate positions bridging the relevant host clusters (Supplementary Figure 3b).

Beyond the global embedding structure, we examined the model’s internal attention patterns to identify which genomic features drive host discrimination. The attention pooling layer assigns a learned weight to each nucleotide position. For each sequence in the host-restricted test dataset, we extracted the top 100 most-attended positions and computed their distribution across the eight genome segments (detailed in Methods). The attention patterns differed markedly among hosts: human-hosted sequences showed concentrated attention in HA and PB2, swine-hosted sequences in HA and NA, and avian-hosted sequences in MP and NP (Figure 3c). Notably, the prominence of HA, PB2, and NP is biologically meaningful, as these segments have been experimentally demonstrated to influence host tropism in influenza A virus[21, 22].

To gain site-resolution insights, we focused on the PB2 segment, given its well-established role in host tropism. We aligned all host-restricted test sequences to a PB2 reference sequence (detailed in Methods) and extracted the per-position attention weights. Among the top 100 positions most attended by the model, 15 sites lie within 10 bp of experimentally reported host adaptation functional sites (Supplementary Table 1), indicating that the model captures biologically relevant sequence features underlying viral host tropism. Moreover, the model exhibits host-specific differentiation in how it weighs these positions. For example, position 661, a well-established host-range determinant amino acid position[7], receives substantially higher attention weight in human-hosted PB2 sequences than the global average computed across the entire test dataset (Figure 3d, 3e, Supplementary Figure 4). Extending this analysis to HA and NP revealed consistent patterns: the sites prioritized by the model overlapped with the known host-tropism-associated amino acid sites, including HA86, HA129, HA134, NP33, and NP53[23] (Supplementary Figure 5, Supplementary Figure 6, Supplementary Table 1).

Together, these results demonstrate that InfluProto not only distinguishes hosts at the sequence level but also assigns disproportionately high attention to biologically relevant functional sites in a host-specific manner, reinforcing the biological plausibility of its learned representations.

### Prototype Distance Quantifies the Continuum of Host Adaptation

We define the cosine distance between a viral sequence and a host-specific prototype as the *host adaptation distance*, which quantitatively characterizes the proximity of the virus to the corresponding host-adapted state in the learned embedding space. To determine whether these distances capture the continuous nature of host adaptation, we performed four complementary analyses.

#### Prototype separation in embedding space

We first examined the pairwise distances among the three learnable prototypes to assess whether they encoded distinct host-specific representations. All three prototypes showed broadly comparable pairwise distances, confirming that they were sufficiently separated to distinguish the three host groups (Supplementary Figure 7). Notably, the human–swine and swine–avian prototype distances were both smaller than the human–avian distance, a pattern consistent with the recognized role of swine as a mixing vessel that facilitates interactions between avian- and human-adapted viruses.

#### Host adaptation distance discriminates host groups and zoonotic signals

We next calculated host adaptation distances for each viral sequence across the four datasets and averaged them within each dataset. Figure 4a presents the resulting distance profiles as a heatmap, with sequences grouped by dataset and sampling host. For the host-restricted test dataset, sequences consistently showed the shortest mean distance to the prototype matching their own host species. This host-specific pattern was also evident within the pdm09 lineage: human-sampled pdm09 viruses were closest to the human prototype, whereas swine-sampled pdm09 viruses were closest to the swine prototype, indicating that InfluProto captures biologically meaningful variation in host adaptation even within this dual-host lineage. Notably, regardless of sampling host, pdm09 viruses exhibited an intermediate mean human adaptation distance—lying between those of human- and swine-hosted viruses—consistent with an intermediate host-adaptation state. The same symmetric pattern was observed for zoonotic avian and zoonotic swine viruses (Figure 4a), collectively demonstrating that host adaptation distance effectively captures the dual-host signatures characteristic of zoonotic viruses.

**Figure 4.**
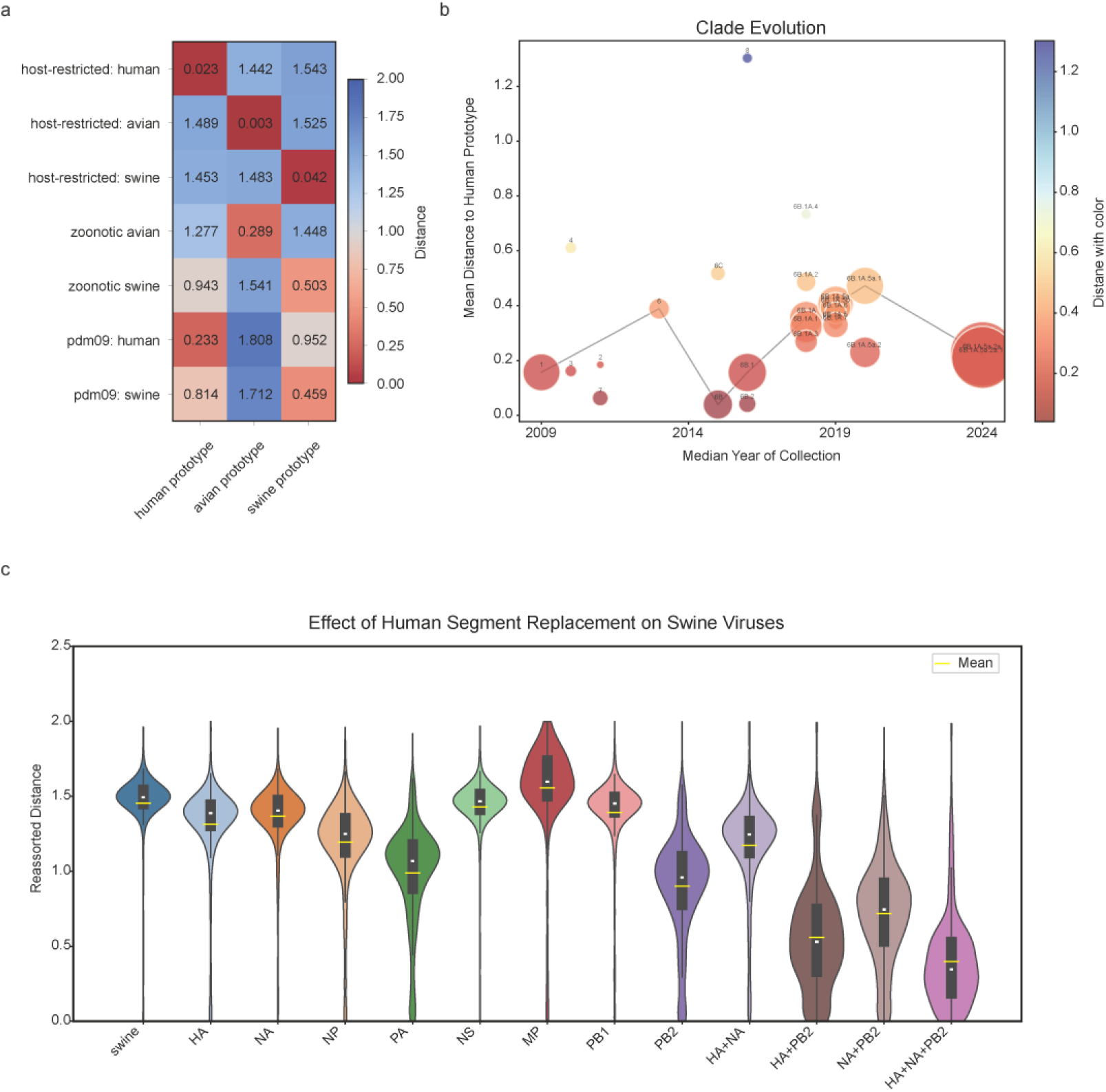
Host adaptation distance as a metric of cross-species transmission potential. **a**, Heatmap of mean host adaptation distances to the three host prototypes across four datasets: host-restricted test dataset, *pdm09 lineage dataset*, *zoonotic avian dataset*, and *zoonotic swine dataset*. **b**, Temporal succession of pdm09 clades and their mean human host adaptation distance. Bubble size represents the number of viral sequences, with larger bubbles indicating more sequences. **c**, Effects of segment replacement on the human host adaptation distance of swine influenza viruses.

#### Temporal dynamics of host adaptation distance track viral evolution

We then examined whether *host adaptation distance* can track evolutionary dynamics over time. Using the whole *pdm09 lineage dataset* of almost 90,000 sequences, we re-annotated clade assignments via the Nextstrain[24] pipeline (details in Methods) and computed the human host adaptation distance for each clade (Supplementary Figure 8a).

Figure 4b traces changes in human host adaptation distance across pdm09 clades over time, revealing a wave-like pattern of alternating increases and decreases across successive clades. Clade 1, which predominated early in the observed period, exhibited a relatively small mean distance to the human prototype, whereas clade 6 showed an increased distance as it became predominant. The subsequent emergence of clade 6B coincided with a decrease in distance, followed by alternating fluctuations among its descendant clades. This temporal pattern is consistent with previously reported changes in human adaptation and immune escape during pdm09 evolution[25–27].

Notably, clade 6B.1A.5a.2 was closer to the human prototype than clade 6B.1A.5a.1 (Figure 4b), even though 6B.1A.5a.1 historically predominated in global circulation, likely due to its greater replication fitness [25]. Nevertheless, the subsequent evolution of 6B.1A.5a.2 aligned with the host affinity captured by the distance metric: its descendant subclades, 6B.1A.5a.2a and 6B.1A.5a.2a.1, remained close to the human prototype (Figure 4b), and later acquired enhanced replication fitness[28] and became globally predominant. This trajectory suggests that distance to prototypes can reveal latent zoonotic and pandemic potential.

#### Reassortment shifts host adaptation distance in a synergistic manner

Finally, we assessed whether *host adaptation distance* could capture shifts in host adaptation following reassortment, a typical mechanism of influenza virus evolution (pdm09 itself being a classic triple-reassortment virus). To simulate this process in silico, we computationally replaced individual segments of swine influenza viruses from the host-restricted test dataset with corresponding segments from randomly selected human-restricted influenza viruses (detailed in Methods). Replacing swine-origin or avian-origin segments with human-origin counterparts progressively reduced the human host adaptation distance (Figure 4c, Supplementary Figure 8b, Supplementary Table 2), demonstrating that the model captures this reassortment process and anticipates the resulting shift in host tropism. Furthermore, the relationship was non-linear: the distance reduction from concurrently replacing both HA and PB2 exceeded the sum of reductions from replacing each segment individually (Figure 4c). This non-additivity indicates that the model has learned synergistic inter-segment signals relevant to host adaptation, rather than simply relying on sequence-level similarity to assign distances. Together, these segment replacement results further support the utility of host adaptation distance as a biologically grounded metric for evaluating cross-species transmission risk.

### InfluProto: An Interactive Web Service for Host Adaptation Assessment

To facilitate public access to InfluProto, we deployed an interactive web service. Users can submit viral sequences through two modes on the web interface: pasting sequences directly into a text area or uploading a FASTA file. For each submitted sequence, the service returns the predicted host and distances to all three prototypes. An example file is provided for users to test the service without preparing their own data (Supplementary Figure 9). The web interface supports both English and Chinese. The model weights and all inference code are publicly available to enable local deployment.

## Discussion

In this study, we presented InfluProto, a deep learning model for influenza A virus host prediction that departs from the conventional classification paradigm in favor of prototype learning. By representing host adaptation as a continuous process by calculating cosine distance to learnable prototypes, our approach offers a way to bridge the fundamental mismatch between standard classification methods and the evolutionary nature of influenza viruses. Rather than forcing viral sequences into discrete host categories, prototype learning allows the model to capture the continuous spectrum of host adaptation, and the resulting embedding space naturally reflects the biological reality that zoonotic viruses occupy intermediate positions between host-adapted populations.

The performance gains over the classification baseline were particularly pronounced for zoonotic sequences. The pdm09 lineage illustrates this clearly. Classification models trained with discrete labels tend to collapse the dual-host signal of pdm09 into a single prediction (predominantly swine), whereas InfluProto preserves a nuanced distance profile that reflects the lineage’s true dual-host nature.

The model captures biologically meaningful signatures of host adaptation. The embedding space naturally separates viruses by host, with zoonotic strains falling at intermediate positions between host-restricted clusters. The attention mechanism converges on segments and sites with established roles in host tropism and the model exhibits host-specific differentiation at functionally critical positions such as residue 661 in PB2.

Beyond host prediction accuracy, the distance to prototypes provides a quantitative metric for assessing cross-species transmission and pandemic potential of viruses. Through four complementary analyses, we showed that this metric captures biologically meaningful signals. Host-restricted viruses consistently showed the shortest distance to the prototype corresponding to their sampling host, whereas zoonotic strains exhibited cross-host distance profiles reflecting adaptation signals associated with multiple hosts. Temporal variation in prototype distances of different clades further reflected evolutionary changes in host adaptation. The observed distance patterns may reflect a trade-off between host adaptation and immune escape. The segment replacement experiment further demonstrates that the model can detect simulated reassortment events, highlighting its potential for the early identification of strains with zoonotic risk. Importantly, the non-linear effects observed in joint segment replacements suggest that the model has learned inter-segment interactions, rather than merely aggregating independent segment-level features.

Several limitations of the present study should be noted. First, the model is restricted to only three main host types: human, swine and avian, without considering rare types. Second, while distance to prototypes correlates with known zoonotic phenotypes, the metric has not yet been independently validated through wet-laboratory validation. Therefore, the metric should be regarded as a computational risk indicator rather than a direct measure of biological infectivity. Third, we only explored cosine distance in this study. The choice of distance function has a substantial impact on prototype learning, and other distance formulations may better capture the biological characteristics of viral evolution. More broadly, representation learning strategies beyond prototype-based learning warrant further investigation to better capture the continuous nature of host adaptation. Fourth, the model was trained exclusively with host-of-isolation as the supervisory signal. However, the pandemic potential of a viral strain is governed by multiple factors beyond host tropism, including replication fitness, immune escape, and transmissibility. Incorporating additional supervisory signals through multi-task learning or auxiliary objectives may improve the model’s ability to assess both cross-species transmission potential and pandemic risk.

Looking ahead, several promising directions warrant further exploration. In principle, the framework could be extended to other viruses characterized by multi-host ecology and sufficient sequence data. The prototype-based architecture is computationally lightweight and readily extensible: a new host can be incorporated by adding a corresponding prototype vector, without redesigning the overall model architecture. As globalization and environmental disruption accelerate the emergence of zoonotic diseases, computational tools that can quantify and monitor cross-species transmission risk will be an increasingly important component of global health surveillance.

## Methods

### Model Architecture

InfluProto consists of three components: a MEGA encoder, an attention pooling layer, and a prototype learning head.

The input comprises the sequences of all eight segments of an influenza A virus, which are concatenated into a single sequence with a “ <sep>” token inserted between adjacent segments. The resulting sequence is then tokenized using a custom tokenizer with a vocabulary of 11 tokens.

The backbone is a MEGA encoder with two MEGA blocks. The encoder outputs per-position embeddings of shape *B* × *L* × 128, where *B* is the batch size and *L* is the sequence length. An attention pooling layer then computes a scalar importance score for each position via a linear projection, normalizes these scores across all positions with a SoftMax operation:

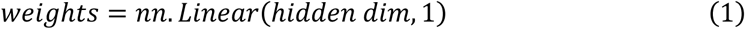

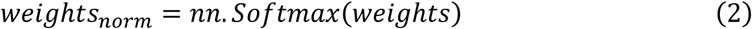

The normalized weights are then used to compute a weighted sum of the position-level embeddings, yielding a single 128-dimensional embedding vector (E(v)) for each viral sequence.

The prototype learning head contains three learnable prototype vectors, *P*_ℎ_, *P_s_*, and *P_a_*, corresponding to human, swine, and avian hosts, respectively. Each prototype has 128 dimensions and is jointly optimized with the encoder parameters during training. For each viral sequence, the distance to each prototype is computed as:

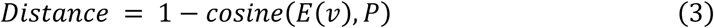

We adjust the cosine similarity to make it more intuitive: the shorter the distance, the closer the pathogen is to a particular prototype, and the more likely it is to infect that host.

The negative distances between each viral embedding and the three host prototypes were used as logits and converted into class probabilities using the softmax function. Cross-entropy loss was then computed using the sampling host of each virus as the ground-truth label:

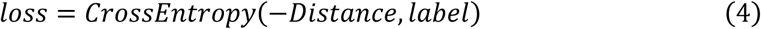

The model was trained with a batch size of 4, a learning rate of 2 × 10⁻⁵, for 70 epochs using the AdamW optimizer.

For comparison, we constructed two baseline classification models using the same MEGA encoder backbone. Baseline 1 retained the attention pooling module but replaced the prototype learning head with a linear classifier. Baseline 2 replaced the attention pooling module with mean pooling and used the same linear classifier. All models were trained using identical optimization settings.

### Datasets

All sequences were obtained from GISAID as of January 5, 2026. Viral segments belonging to the same isolate were grouped by Isolate ID and only isolates with complete full-genome sequences were retained. Duplicate sequences were removed by hash-based deduplication. Host metadata were mapped to three categories: human, swine, and avian. Sequences with other or ambiguous host annotations were excluded. (1) *2009 pandemic H1N1(pdm09) lineage dataset*: sequences whose lineage annotation contained “pdm09” were extracted and reserved for independent evaluation. (2) *Zoonotic avian dataset*: sequences with pathogenicity annotated as LPAI or HPAI but sampled from a human host. (3) *Zoonotic swine dataset*: sequences with swine-origin metadata notes but sampled from a human host. (4) *Host-restricted dataset*: approximately 30,000 sequences were sampled and partitioned into training, validation, and test sets at an 8:1:1 ratio. Sequences belonging to the *pdm09 lineage dataset*, *zoonotic avian dataset*, and *zoonotic swine dataset* were excluded to ensure unambiguous host-restricted labels during training. The curated collection was used to construct four datasets.

### Identification of Important Segments and Sites

To identify the genomic regions prioritized by the model, we extracted the 100 positions with the highest attention weights for each sequence in the host-restricted test dataset. Segment boundaries were identified by the positions of “<sep>” tokens, and the distribution of the top 100 positions across the eight segments was computed separately for each host category.

For site-resolution analysis, we focused on the PB2 segment. Since viral sequences vary in length, all PB2 sequences from the host-restricted test dataset were first aligned to a reference PB2 sequence using MMSeqs2[23] multiple sequence alignment. After alignment, the per-position attention weights from each sequence were mapped to the common reference coordinate system. This allowed us to compute the mean attention weight at each position and to compare these profiles against experimentally reported functional sites mapped to the same coordinates. Analyses of HA and NP followed the same procedure.

### Clade reannotation

Because the definitions of pdm09 clades have changed over time, the original GISAID clade annotations may not be directly comparable across different collection periods. To obtain consistent clade assignments, we reannotated all sequences in the *pdm09 lineage dataset* based on their HA sequences using the Nextstrain pipeline and the flu_h1n1pdm_ha reference. The mean distance to the human prototype was then calculated for each clade by averaging the distances of all viral sequences within that clade to the human prototype. To trace how the distance to the human prototype changed over time, we considered only clades comprising more than 1,000 sequences. When multiple clades had overlapping collection periods, the clade containing the largest number of sequences was selected as the representative to reduce redundancy.

### Segment Replacement

To simulate segment reassortment in silico, we computationally replaced individual segments of swine influenza viruses in the host-restricted test dataset with the corresponding segments from randomly selected human influenza viruses. The replacement was performed at the sequence level: the original segment was substituted with the human-origin counterpart, and the resulting eight-segment sequence was fed to the model. Single-segment replacement was performed for each of the eight segments independently. For multi-segment replacement, the same procedure was applied concurrently to selected segment pairs. The distance to the human prototype was computed for each original and segment-replaced sequence.

## Data Availability

The GISAID accession numbers for all sequences used in this study are available at Zenodo (https://doi.org/10.5281/zenodo.21234825). Individual sequence data can be obtained from the GISAID EpiFlu database (https://www.gisaid.org) upon registration.

## Code Availability

The source code for InfluProto is publicly available at https://github.com/GZZHY79/InfluProto under the MIT license. The repository includes the model architecture, a custom tokenizer, inference scripts, an example input file, and a complete set of dependencies (requirements.txt). An online prediction tool is also available at https://ngdc.cncb.ac.cn/bit/influProto, providing a web-based interface for sequence submission and prediction.

## Acknowledgements

This work was supported by the National Key Research & Development Program of China (2024YFC2311303), the National Natural Science Foundation of China (32270718), the National Key Research & Development Program of China (2025YFF1207901, 2023YFC2604400).

## Author Contributions

Z.G. and S.S. conceptualized the study. Z.G. and L.L. designed the experiments. Z.G. conducted the experiments and analyzed the results. Z.G. and L.L. wrote the original draft. R.Y. and Y.W. contributed valuable discussions and scientific advice throughout the study. S.S. supervised the study and critically revised the manuscript. All authors reviewed and approved the final manuscript.

## References

1 Rahman, M. T. et al. Zoonotic Diseases: Etiology, Impact, and Control. Microorganisms 8 (2020). 10.3390/microorganisms8091405

2 Bernstein, A. S. et al. The costs and benefits of primary prevention of zoonotic pandemics. Science Advances 8, eabl4183 (2022). doi:10.1126/sciadv.abl4183

3 Abdelwhab, E. M. & Mettenleiter, T. C. Zoonotic Animal Influenza Virus and Potential Mixing Vessel Hosts. Viruses 15 (2023). 10.3390/v15040980

4 Johnson, K. E. E., Song, T., Greenbaum, B. & Ghedin, E. Getting the flu: 5 key facts about influenza virus evolution. PLOS Pathogens 13, e1006450 (2017). 10.1371/journal.ppat.1006450

5 Belser, J. A. et al. Robustness of the Ferret Model for Influenza Risk Assessment Studies: a Cross-Laboratory Exercise. mBio 13, e0117422 (2022). 10.1128/mbio.01174-22

6 Nguyen, T. Q., Rollon, R. & Choi, Y. K. Animal Models for Influenza Research: Strengths and Weaknesses. Viruses 13 (2021). 10.3390/v13061011

7 Lu, C. et al. FluPhenotype—a one-stop platform for early warnings of the influenza A virus. Bioinformatics 36, 3251–3253 (2020). 10.1093/bioinformatics/btaa083

8 Livieratos, A., Kagadis, G. C., Gogos, C. & Akinosoglou, K. AI Methods Tailored to Influenza, RSV, HIV, and SARS-CoV-2: A Focused Review. Pathogens 14 (2025). 10.3390/pathogens14080748

9 Ye, R., Li, L., Ribeiro de Vasconcelos, A. T. & Song, S. Influ-BERT: a domain-adaptive genomic language model for advancing influenza A virus research. Briefings in Bioinformatics 27 (2026). 10.1093/bib/bbag171

10 Kieran, T. J., Sun, X., Maines, T. R. & Belser, J. A. Machine learning approaches for influenza A virus risk assessment identifies predictive correlates using ferret model in vivo data. Communications Biology 7, 927 (2024). 10.1038/s42003-024-06629-0

11 Shang, J. & Sun, Y. CHERRY: a Computational metHod for accuratE pRediction of virus-pRokarYotic interactions using a graph encoder-decoder model. Brief Bioinform 23 (2022). 10.1093/bib/bbac182

12 Mock, F., Viehweger, A., Barth, E. & Marz, M. VIDHOP, viral host prediction with deep learning. Bioinformatics 37, 318–325 (2021). 10.1093/bioinformatics/btaa705

13 Hu, M. et al. Flu-CNN: identifying host specificity of Influenza A virus using convolutional networks. Hum Genomics 19, 96 (2025). 10.1186/s40246-025-00812-y

14 Hatibi, N. et al. Misclassified: identification of zoonotic transition biomarker candidates for influenza A viruses using deep neural network. Frontiers in Genetics **Volume** 14 - 2023 (2023). 10.3389/fgene.2023.1145166

15 Yang, J. et al. Using Artificial Intelligence to Assess Cross-Species Transmission Potential of Influenza A Virus. bioRxiv, 2025.2006.2017.660059 (2025). 10.1101/2025.06.17.660059

16 Snell, J., Swersky, K. & Zemel, R. S. in Neural Information Processing Systems.

17 Vinyals, O., Blundell, C., Lillicrap, T. P., Kavukcuoglu, K. & Wierstra, D. in Neural Information Processing Systems.

18 Khare, S. et al. GISAID’s Role in Pandemic Response. China CDC Weekly 3, 1049–1051 (2021). 10.46234/ccdcw2021.255

19 Potdar, V. A. et al. Genetic characterization of the influenza A pandemic (H1N1) 2009 virus isolates from India. PLoS One 5, e9693 (2010). 10.1371/journal.pone.0009693

20. Xuezhe, M., et al. in The Eleventh International Conference on Learning Representations (2023).

21 Ciminski, K., Chase, G. P., Beer, M. & Schwemmle, M. Influenza A Viruses: Understanding Human Host Determinants. Trends in Molecular Medicine 27, 104–112 (2021). 10.1016/j.molmed.2020.09.014

22 Mänz, B. et al. Pandemic Influenza A Viruses Escape from Restriction by Human MxA through Adaptive Mutations in the Nucleoprotein. PLOS Pathogens 9, e1003279 (2013). 10.1371/journal.ppat.1003279

23 Steinegger, M. & Söding, J. MMseqs2 enables sensitive protein sequence searching for the analysis of massive data sets. Nature Biotechnology 35, 1026–1028 (2017). 10.1038/nbt.3988

24 Hadfield, J. et al. Nextstrain: real-time tracking of pathogen evolution. Bioinformatics 34, 4121–4123 (2018). 10.1093/bioinformatics/bty407

25 Swanson, N. J. et al. 2019-2020 H1N1 clade A5a.1 viruses have better in vitro fitness compared with the co-circulating A5a.2 clade. Sci Rep 13, 10223 (2023). 10.1038/s41598-023-37122-z

26 de Vries, R. P. et al. Evolution of the hemagglutinin protein of the new pandemic H1N1 influenza virus: maintaining optimal receptor binding by compensatory substitutions. J Virol 87, 13868–13877 (2013). 10.1128/jvi.01955-13

27 Su, Y. C. F. et al. Phylodynamics of H1N1/2009 influenza reveals the transition from host adaptation to immune-driven selection. Nat Commun 6, 7952 (2015). 10.1038/ncomms8952

28 Swanson, N. *CHARACTERIZATION OF IN VITRO FITNESS AND ANTIGENIC* DRIFT DIFFERENCES BETWEEN CIRCULATING H1N1 INFLUENZA CLADES, Johns Hopkins University, (2024).

