## Supplementary Information for "InfluProto: Predicting Cross-Species Transmission of Influenza A Virus via Prototype Learning"

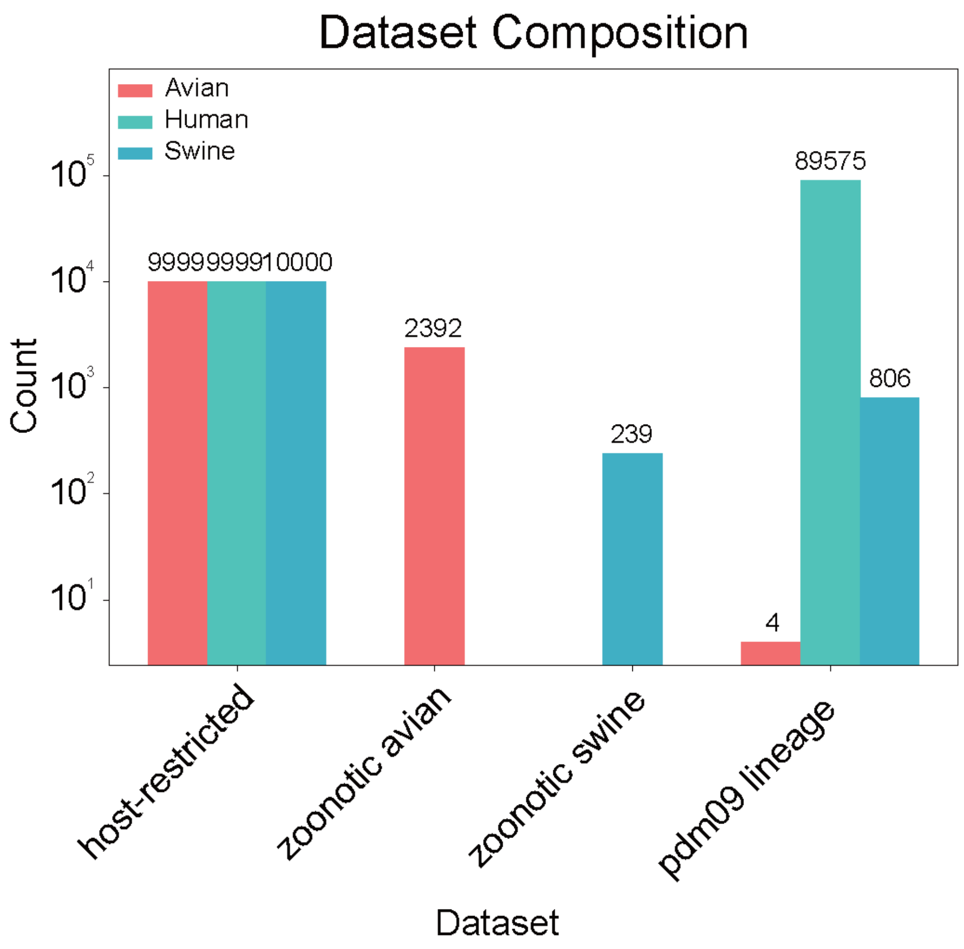


**Supplementary Figure 1. Datasets Composition.**

Bar chart showing the number of sequences in each dataset: h*ost-restricted dataset*, *zoonotic avian dataset*, *zoonotic swine dataset*, and *pdm09 dataset*.

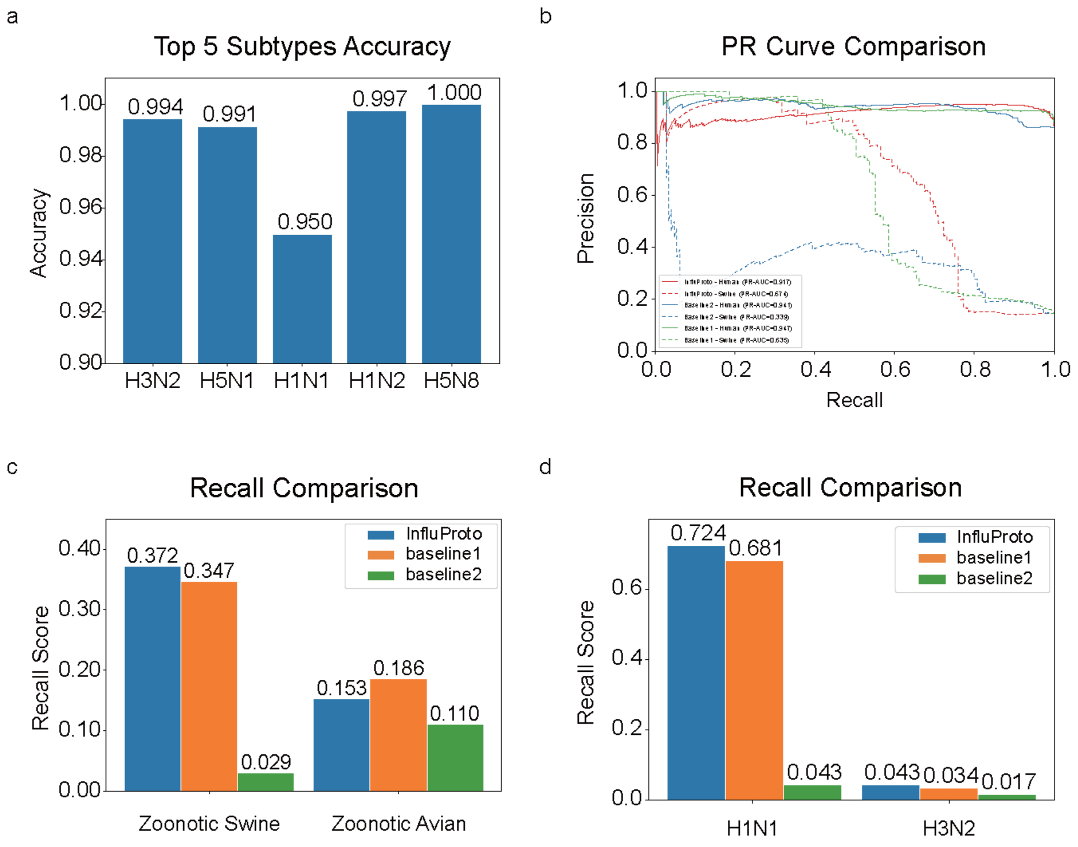


#### Supplementary Figure 2 Additional performance metrics.

**a.** Prediction accuracy for the top five most prevalent subtypes in the host restricted test dataset.

**b.** Precision–recall curves of InfluProto, baseline model 1, and baseline model 2 across different virus categories. Solid and dashed lines distinguish different virus types; models sharing the same color belong to the same model family. InfluProto is shown in red, baseline model 1 in blue, and baseline model 2 in green.

**c.** Recall comparison of InfluProto and baseline models on two zoonotic virus datasets: *zoonotic swine dataset* and *zoonotic avian dataset*. InfluProto achieves the highest recall on *zoonotic swine dataset* and outperforms baseline model 1 on *zoonotic avian dataset*.

**d.** Recall comparison by viral subtype within the *zoonotic swine dataset*. InfluProto shows the highest recall among the three models across all tested subtypes.

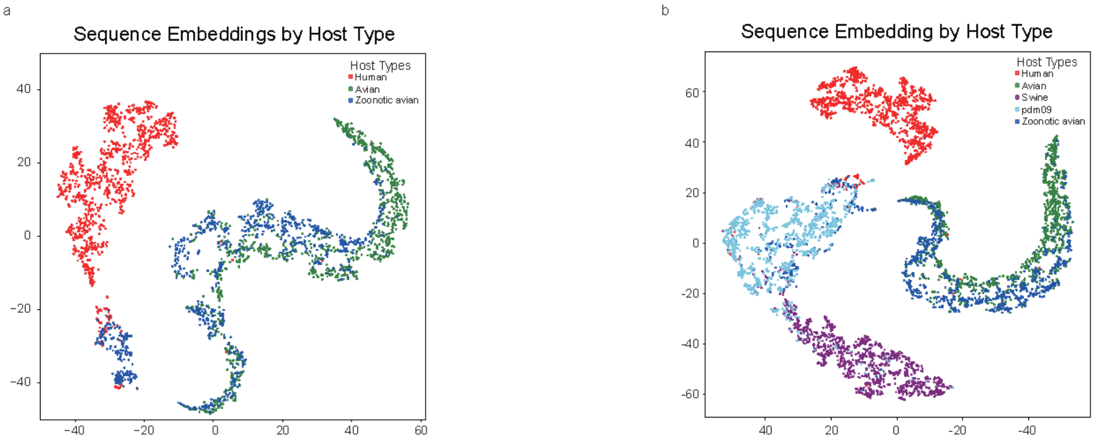


**Supplementary Figure 3. Additional t-SNE embedding visualizations.**

- **a.** t-SNE projection of viral embeddings from three host types: host-restricted human, host-restricted avian, and zoonotic avian, colored by category.
- **b.** t-SNE projection of viral embeddings from five virus categories, colored by category.

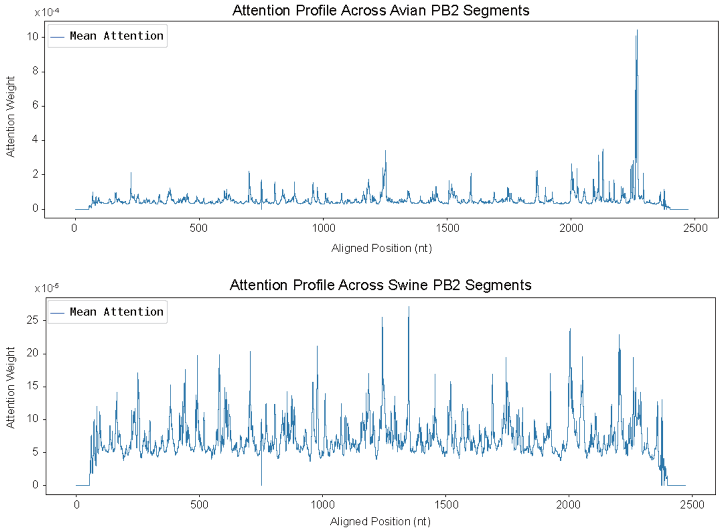


### Supplementary Figure 4. Per-position attention profiles across the PB2 segment.

Attention weights of InfluProto mapped to the PB2 reference sequence, shown separately for avian derived viruses and swine derived viruses.

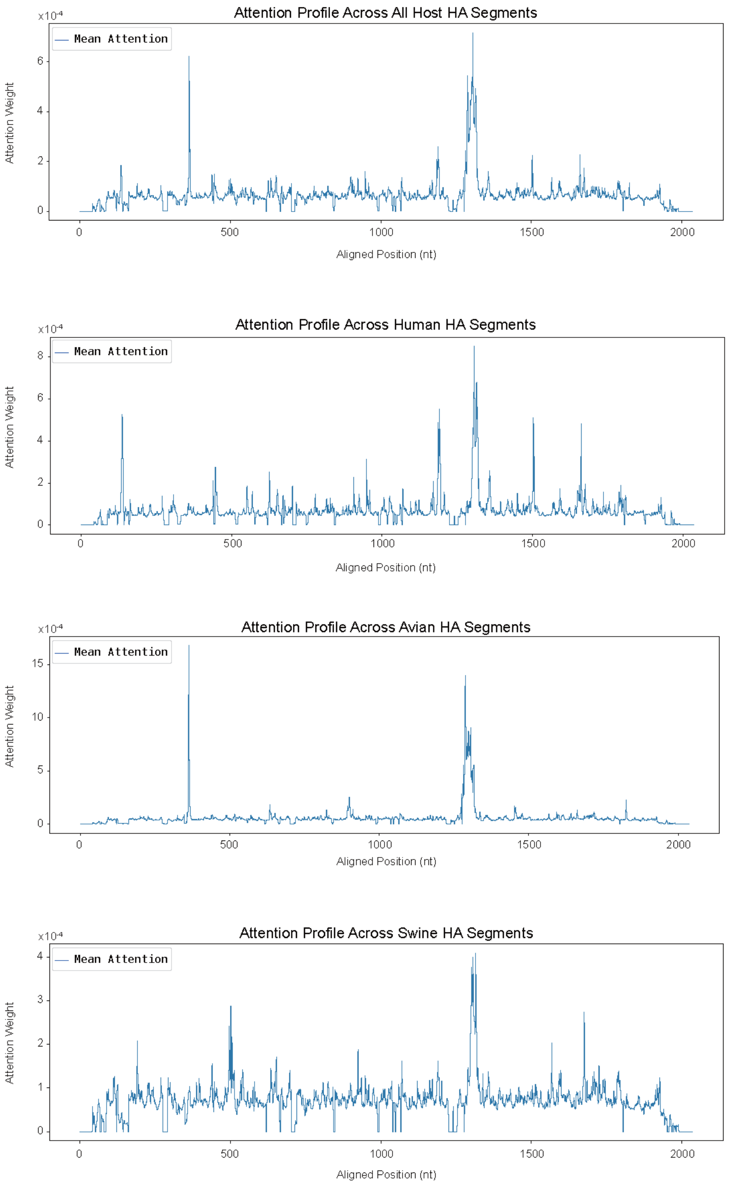


#### Supplementary Figure 5. Per-position attention profiles across the HA segment.

Attention weights of InfluProto mapped to the HA reference sequence, shown separately for all viruses (global), human derived viruses, avian derived viruses, and swine derived viruses.

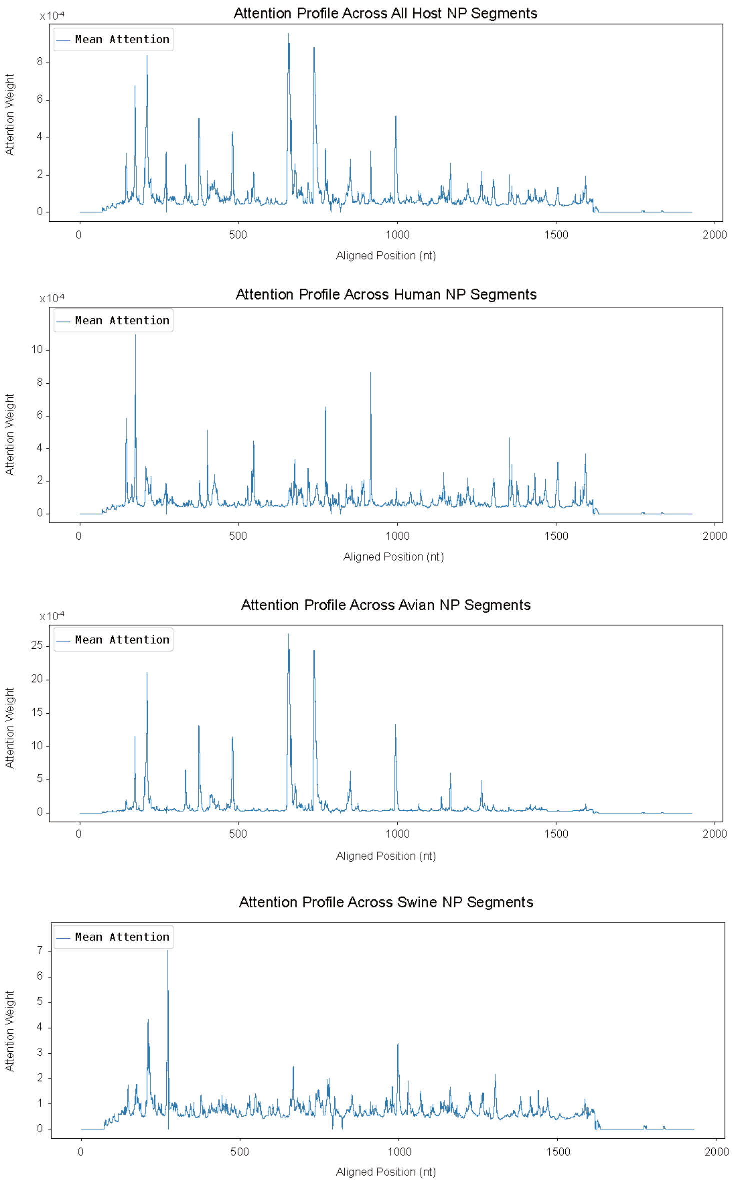


#### Supplementary Figure 6. Per-position attention profiles across the HA segment.

Attention weights of InfluProto mapped to the NP reference sequence, shown separately for all viruses (global), human derived viruses, avian derived viruses, and swine derived viruses.

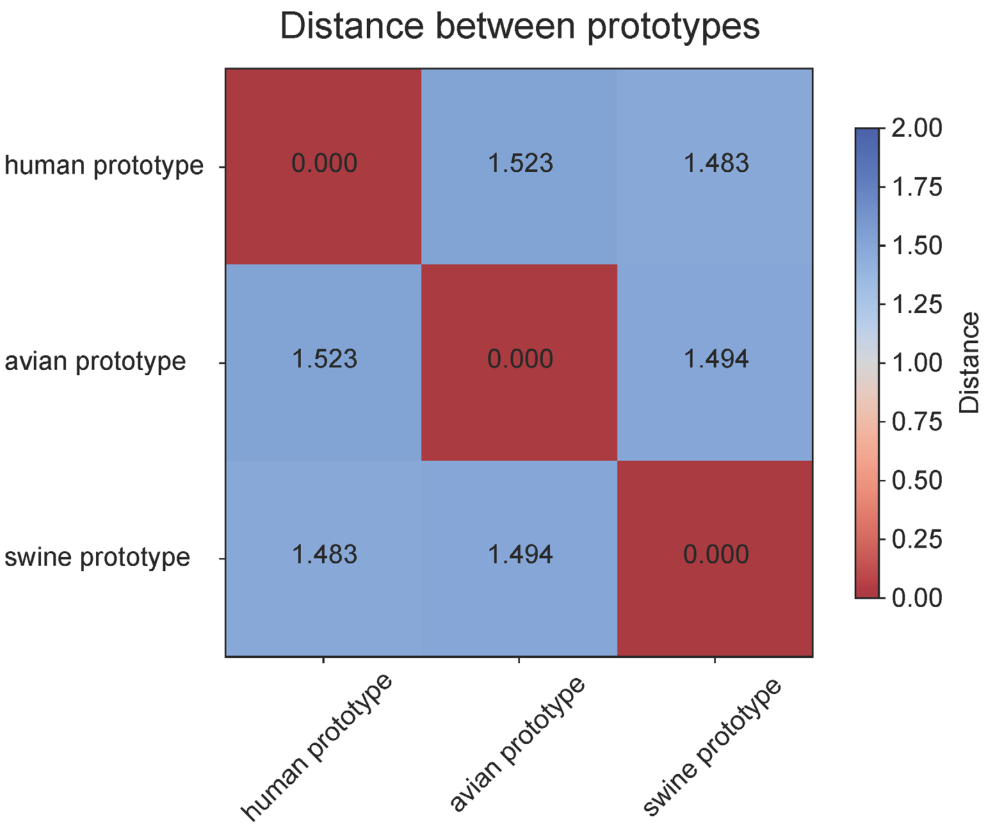


**Supplementary Figure 7. Pairwise distances between learned prototype vectors.**

Heatmap of distances between the three learned prototypes. Redder color indicates shorter distance meaning greater similarity between prototypes.

a

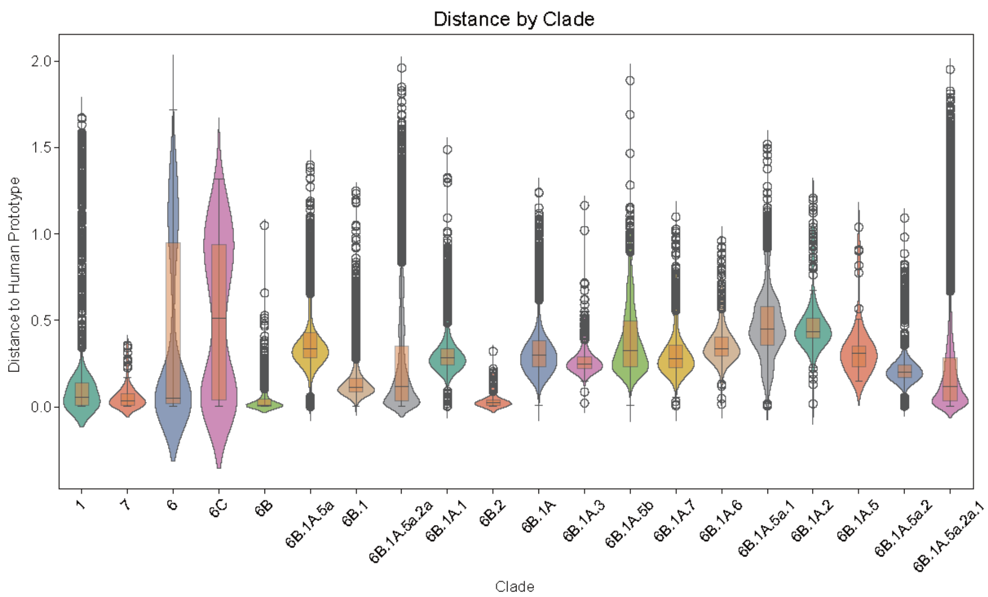


b

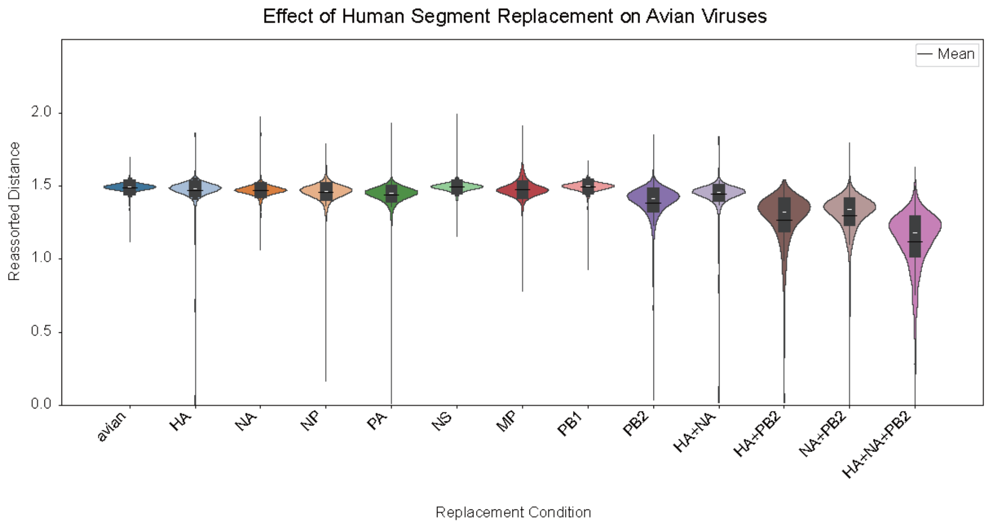


#### Supplementary Figure 8. Additional distance-to-prototype analyses.

**a.** Violin plots show the distribution of distances to the human prototype across pdm09 clades. Each violin corresponds to one clade.

**b.** Mean distance to the human prototype before and after individual and multi-segment replacement for each of the eight genome segments of avian-hosted virus.

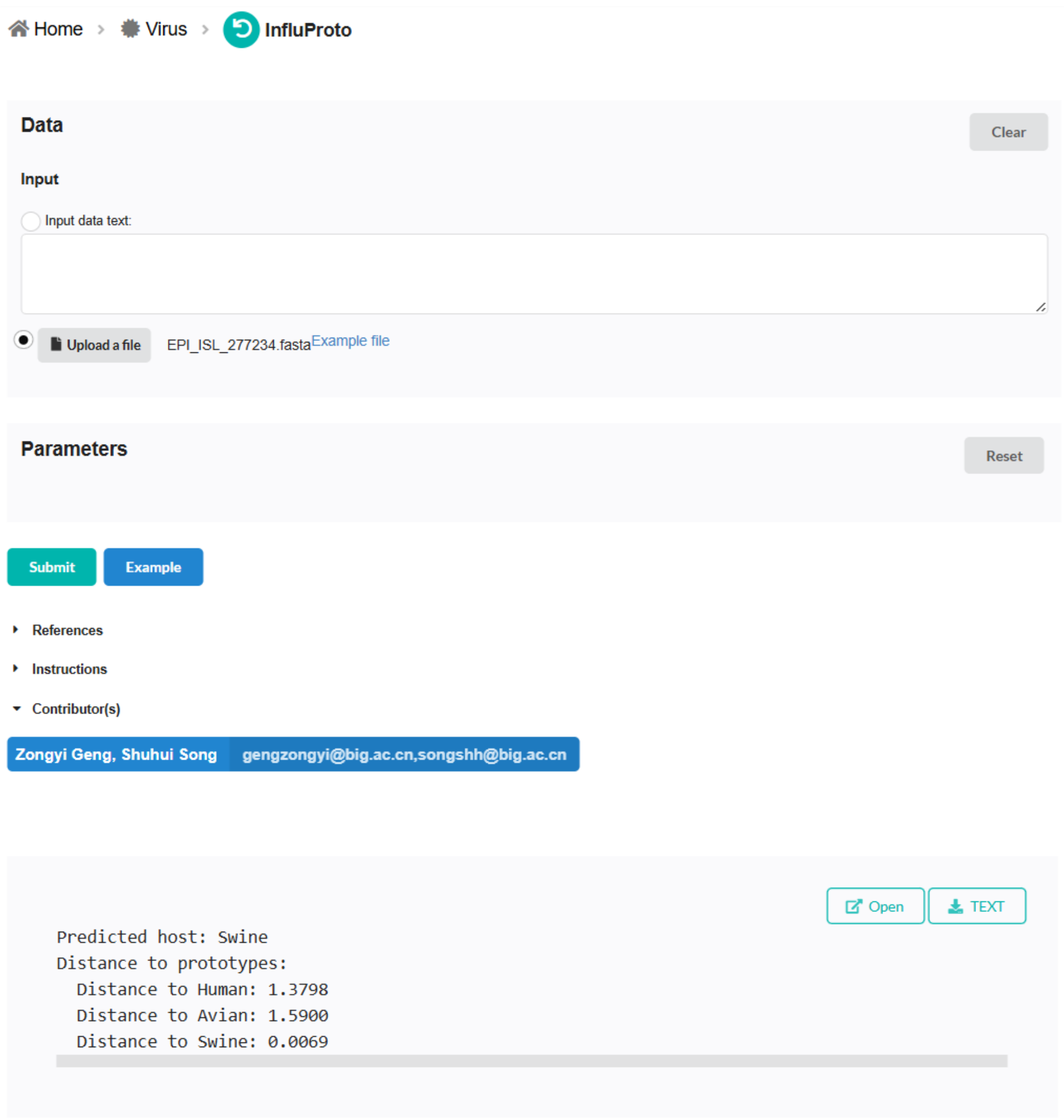


#### Supplementary Figure 9. Graphic web service of InfluProto.

The web service supports two ways to submit influenza A virus sequence. After calculating, the web service returns the predicted host and distances to all three prototypes.

| Segment | Class | Position | Number |
|---|---|---|---|
| PB2 | Reported sites | 44, 64, 191, 199, 256, 271, 323, 355, 391, 447, 475, 477, 504, 529, 567, 588, 590, 613, 627, 636, 661, 674, 701, 714 | 24 |
| | Corresponding nucleotide sites | 213, 273, 654, 678, 850, 895, 1051, 1147, 1255, 1423, 1507, 1513, 1594, 1669, 1783, 1846, 1852, 1921, 1963, 1990, 2065, 2104, 2185, 2224 | 24 |
| | All viruses' attention sites | 1245, 1247, 1248, 1249, 1250, 1251, 1252, 1253, 1257, 1520, 1925, 2055, 2110, 2111, 2112 | 15 |
| | Human hosted viruses' attention sites | 1257, 1258, 1925, 2055, 2109, 2110, 2111 | 7 |
| | Swine hosted viruses' attention sites | 885, 1245, 1246, 1247, 1248, 1249, 1250, 1251, 1252, 1253, 1254, 1508, 1509, 1520, 1596, 1597, 1598, 1860, 1861, 2096, 2111, 2112, 2113, 2175 | 17 |
| | Avian hosted viruses' attention sites | 855, 856, 1245, 1246, 1247, 1248, 1249, 1250, 1520, 1521, 1924, 1925, 1999, 2055, 2056, 2057, 2058 | 24 |
| HA | Reported sites | 33, 53, 100, 283, 289, 305, 313, 316, 350, 351, 353, 357 | 12 |
| | Corresponding nucleotide sites | 214, 275, 416, 968, 986, 1034, 1058, 1067, 1169, 1172, 1178, 1190 | 12 |
| | All virus attention sites | 206, 207, 208, 209, 210, 211, 212, 213, 214, 215, 222, 224, 271, 272, 993, 994, 995, 996, 1167, 1168 | 20 |
| | Human hosted viruses' attention sites | 207, 208, 209, 210, 211, 212, 213, 214, 215, 224, 271, 272, 420, 422, 423, 424, 425, 426 | 18 |

| | | | |
|---|---|---|---|
| | Swine hosted viruses' attention sites | 206, 207, 208, 209, 210, 211, 212, 213, 214, 215, 216, 217, 218, 267, 268, 269, 270, 271, 272, 274, 275, 276, 960, 979, 994, 995, 996, 1028, 1029, 1068, 1160, 1161, 1162, 1163 | 34 |
| | Avian hosted viruses' attention sites | 204, 205, 206, 207, 208, 209, 210, 211, 212, 213, 214, 215, 411, 414, 415, 416, 417, 993, 994, 995, 996, 1167, 1168, 1169 | 24 |
| NP | Reported sites | 18, 75, 86, 94, 129, 134, 139, 144, 154, 158, 182, 190, 192, 193, 222, 223, 224, 225, 226, 227, 228, 248, 263, 315, 388, 435, 444, 497, 522 | 29 |
| | Corresponding nucleotide sites | 179, 375, 408, 432, 537, 552, 567, 582, 612, 627, 723, 747, 753, 756, 848, 851, 854, 857, 860, 863, 866, 926, 971, 1142, 1389, 1530, 1557, 1716, 1791 | 29 |
| | All viruses' attention sites | 365, 366, 439, 626, 634, 635, 924, 1788 | 8 |
| | Human hosted viruses' attention sites | 439, 550, 551, 552, 569, 626, 924, 1788, 1794 | 9 |
| | Swine hosted viruses' attention sites | 438, 439, 539, 540, 541, 542, 634, 635, 923, 924, 925, 935, 1567, 1708, 1709, 1725, 1786, 1788, 1790, 1791 | 20 |
| | Avian hosted viruses' attention sites | 365, 366, 367, 368, 634, 635, 636, 1718, 1720 | 9 |

**Supplementary Table 1. Host-tropism-associated functional sites and InfluProto attention sites.**

Reported amino acid positions and corresponding nucleotide positions from the literature that are associated with host tropism in PB2, HA, and NP, alongside InfluProto's top-attended nucleotide positions within these segments. Positions are reported globally (all viruses) and stratified by host: human, swine, and avian.

| Host | Segment | P-value | FDR Q-value |
|---|---|---|---|
| Swine | HA | 0.000 | 0.000 |
| | NA | 0.000 | 0.000 |
| | NP | 0.000 | 0.000 |
| | PA | 0.000 | 0.000 |
| | NS | 0.000 | 0.000 |
| | MP | 0.000 | 0.000 |
| | PB1 | 0.000 | 0.000 |
| | PB2 | 0.000 | 0.000 |
| | HA+NA | 0.000 | 0.000 |
| | HA+PB2 | 0.000 | 0.000 |
| | NA+PB2 | 0.000 | 0.000 |
| | HA+NA+PB2 | 0.000 | 0.000 |
| Avian | HA | 0.000 | 0.000 |
| | NA | 0.000 | 0.000 |
| | NP | 0.000 | 0.000 |
| | PA | 0.000 | 0.000 |
| | NS | 0.018 | 0.018 |
| | MP | 0.000 | 0.000 |
| | PB1 | 0.000 | 0.000 |
| | PB2 | 0.000 | 0.000 |
| | HA+NA | 0.000 | 0.000 |
| | HA+PB2 | 0.000 | 0.000 |
| | NA+PB2 | 0.000 | 0.000 |
| | HA+NA+PB2 | 0.000 | 0.000 |

**Supplementary Table 2. Statistical significance of segment replacement distance changes.**

Mean distance to the human prototype after segment replacement, tested against the baseline distance of unreplaced sequences (one-sample t-test). Raw P-values and FDR-corrected Q-values (Benjamini–Hochberg procedure) are reported for each single-segment and multi-segment replacement condition. Tests are performed separately for swine segment-to-human segment and avian segment-to-human segment replacement settings.
